# Constraint and innovation in color evolution among species and among plumage patches in five avian radiations

**DOI:** 10.1101/2023.08.02.551664

**Authors:** Chad M. Eliason, Rafael S. Marcondes, Muir D. Eaton, Rafael Maia, Kevin J. Burns, Allison J. Shultz

## Abstract

Understanding the causes and limits of phenotypic diversification remains a key challenge in evolutionary biology. Color patterns are some of the most diverse phenotypes in nature. In birds, recent work within families has suggested that plumage complexity might be a key innovation driving color diversity. Whether these patterns hold at larger taxonomic scales remains unknown. Here, we assemble a large database of UV-Vis spectral data across five diverse clades of birds (45791 spectra, 1135 species). Using multivariate phylogenetic comparative methods, we compare evolutionary rates and color space occupancy (i.e., quantification of observed colors) among these clades. Novel color-producing mechanisms have enabled clades to occupy new regions of color space, but using more coloration mechanisms did not result in overall more color space occupancy. Instead, the use of more color-producing mechanisms resulted in faster rates of color evolution and less integrated color among plumage regions. Flexible Bayesian modeling further allowed us to assess the relationship between interpatch and interspecific directions of color variation. We find that interpatch variation generally diverges from interspecies cladewise trends in males but not females, suggesting developmental or selective constraints operating in females across evolutionary scales. By comparing rates among clades and assessing both interpatch and interspecies color variation, we reveal how innovations and constraints operate across evolutionary and developmental scales.

## Main

For complex phenotypes made up of two or more traits, adaptive radiation theory predicts that phenotypic evolution should occur along the major axis of an ellipse describing covariation between subtraits–that is, along genetic lines of least resistance^1^. Depending on the direction of selection relative to the major axes of phenotypic variation, strong covariation between subtraits can either reduce or enhance rates of phenotypic evolution^2^. Recent empirical evidence in avian beaks suggests that evolutionary innovations can cause discrete jumps in phenotypic space and/or reorient covariance matrices, with the end result of increasing phenotypic diversity^3^.

Developmental and functional links between subtraits might explain such shifts. For example, in the avian skull, independent developmental trajectories explain evolutionary independence among different regions of the skull^4^. In avian limbs, selection for coordinated function (e.g., flight) can strengthen evolutionary covariation among forelimb and hind limb elements^5^. Despite considerable research on the interplay between development constraint and the evolution of ecological traits^4, 6–9^, less is known about how the evolution of ornamental traits is shaped by developmental or functional constraints^10, 11^. An example of this is color pattern, a multifunctional and frequently ornamental trait highly studied by ecologists and evolutionary biologists, but a trait that is highly multidimensional, making it challenging to study in a comparative framework relative to simpler traits such as limb morphology or wing shape.

Courtship phenotypes of birds, such as birds-of-paradise (Paradisaeidae), comprise some of the most diverse multimodal displays in nature^12^. A major axis of variation in courtship signals is in plumage coloration. Diverse feather colors in birds result from a combination of light scattering by nanostructured feather tissue (keratin, melanin, and air)^13^ and light absorption by pigments deposited in feathers (e.g., melanins, carotenoids)^14, 15^. Innovations in the production and deployment of these different color mechanisms across the body have expanded avian color space^16^. While plumage complexity (i.e., number of distinct colors in a plumage) has been studied extensively in relation to abiotic and biotic factors^12, 17–19^, the role of color patch arrangement in explaining interspecific differences in color patterning has rarely been investigated^20^. Plumage patches are defined by developmentally distinct feather tracts that can vary independently (i.e., they are modular)^21^. A recent study in estrildid finches shows that color maps across the body are spatially conserved among species^22^, suggesting that plumage patch boundaries in a clade remain static while color pattern diversity is determined by downstream differences in regulatory factors that tune the colors of individual patches^23^. Co-expression of feather genes across the body^24^ or expansion of color patches in response to increased pigment gene activity^25^, as has recently been shown in canids, could potentially explain observed evolutionary correlations in color among plumage patches in several clades^20, 26, 27^.

Novel colors in a bird’s plumage can arise either through the evolution of new color-producing mechanisms (e.g., iridescence)^28^ or finescale elaboration of existing mechanisms (e.g., nanoscale changes in feather structure or variation in carotenoid pigment concentrations)^29, 30^. Multifarious selection on multiple developmentally independent color patches would further translate to greater evolutionary potential for divergence in overall color patterns between species^31, 32^. We hypothesized that mechanistic processes (i.e., how and where color is produced on a bird’s body) will explain evolutionary trends in coloration. This hypothesis predicts that i) plumage integration (i.e., the tendency for spatially adjacent patches to display similar colors) and rates of color evolution vary among clades with different color-producing mechanisms due to differences in evolutionary lability; ii) color evolution occurs along the major axis of an ellipse describing covariation between color patches at the individual plumage level (Fig. 1); and iii) clades with a greater number of color-producing mechanisms and lower plumage integration (i.e., more modular plumages) accrue color diversity at a faster rate. Here, we use multivariate comparative methods and flexible Bayesian phylogenetic mixed models to test these predictions using a large spectral data set (46160 reflectance spectra, 8-22 plumage patches per bird) covering 1147 species across five diverse clades–African starlings (Sturnidae), kingfishers (Alcedinidae), tanagers and allies (Thraupidae), blackbirds and allies (Icteridae), and antbirds and ovenbirds (Furnariida). These clades vary in number of color-producing mechanisms and species richness, together encompassing >10% of avian biodiversity^33^. Comparing evolutionary trends among diverse avian clades can shed light on the relative importance of elaboration and innovation in explaining color evolution.

**Fig. 1.**
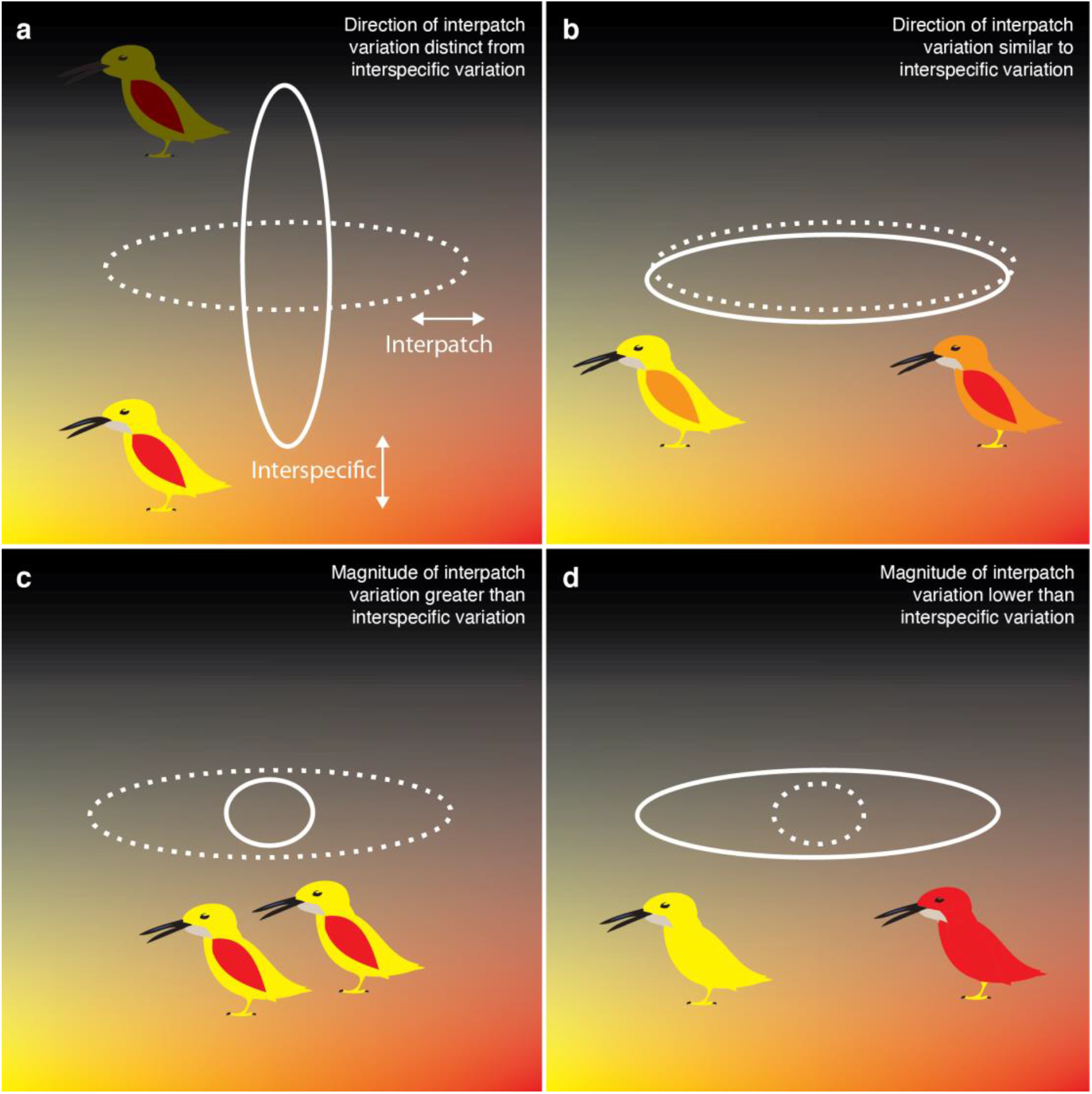
Ways that interpatch and interspecific color variation might differ. Axes depict hypothetical variation in color axes hue (x) and lightness (y). Ellipses show orientation of primary axes of interspecific (solid; between species) and interpatch color variation (dashed lines; between patches - wing versus body). a, species diverge in lightness while patches vary in hue, suggesting decoupling between constraints operating at developmental and interspecific levels. b, species and patches vary along the same axis, suggesting constraints limiting what kinds of colors can be produced across the body are also operating at evolutionary scales. c, birds have complex colorful plumages but there is little variation among species, suggesting weak constraints at the plumage level but evolutionary constraints limiting color evolution. d, color is distributed mainly among species with little interpatch variation, suggesting strong constraints operating at the individual level but diversifying processes operating at evolutionary scales.

## Results

### Birds expand color space using both pigments and nanostructures

We used the avian tetrahedral color space, in which color data are plotted based on the relative stimulation of the four cones in the avian eye^34^, to investigate differences in plumage color within and among clades of birds. To understand differences in color space occupation among the five clades, we compared color space volumes (i.e., the 3-D volume of a convex hull encompassing points in avian color space) in the R package pavo^35^ and classified each patch by overall color-producing mechanism (carotenoids: red, orange, yellow; melanin: phaeomelanin, eumelanin; structural color: structural barb rami, structural barbule (iridescence); white). The estimated color space volume for Thraupidae was almost 5 times larger than any other clade (color space volume = 0.066), while the smallest color space volume was seen in Furnariida (color space volume = 0.012; see Table S1). The color space volume for Icteridae (color space volume = 0.016) was approximately the same as in Alcedinidae (color space volume = 0.015; Fig. 2), but each of these two clades expands color space in different ways, using either yellow and red carotenoids (Icteridae; Fig. 2d) or phaeomelanins and barb-based structural coloration (Alcedinidae; Fig. 2c). We observed the greatest color space similarity between Alcedinidae and Sturnidae (42.6% overlapping, see Table S1), both of which primarily use melanins and structural color, although different types of structural color.

**Fig. 2.**
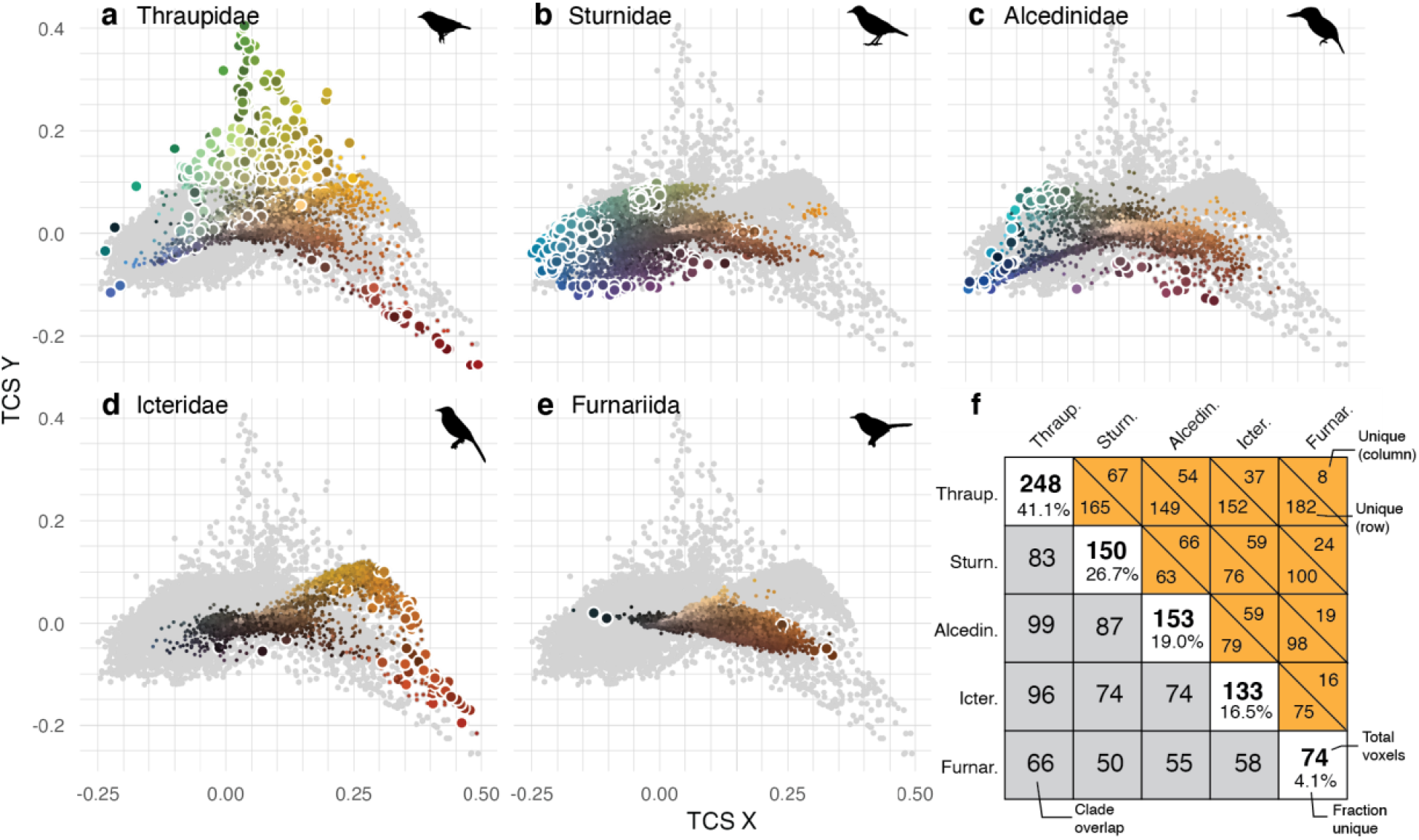
Novelty in color space occupation among clades. Tetrahedral color space plots for (a) Thraupidae, (b) Sturnidae, (c) Alcedinidae, (d) Icteridae, and (e) Furnariida. Gray points are all color space XY coordinates in the data set. Large points encircled in white depict novel colors in a clade. Lines delimit voxels dividing up color space (voxels are the 3-D equivalent of a 2-D pixel in an image). f, color space occupation determined by 0.05 x 0.05 x 0.05 voxels (i.e., grid cells shown in a -e) in tetrahedral color space. See legend for description of numbers in cells.

Voxel-based (the 3-D analog of a 2-D pixel) analysis of color space reveals significant color novelty in several clades, defined as regions of the color space only occupied by that clade. In Thraupidae, mixing of carotenoid yellow pigments and non-iridescent structural blue colors^36^ results in novel green colors within our data set (Figs. 2a, S10). Despite color space overlap between Sturnidae and Alcedinidae, Sturnidae occupies novel regions of color space (e.g., violet and blue colors with low UV content; Fig. S2b) owing to the presence of thin film and multilayer feather structures that produce the characteristic iridescent colors of the clade^28, 37^. Further novelty is seen in Alcedinidae, as mixing of phaeomelanin pigments and structural color mechanisms produce violet colors approaching those seen in Sturnidae, yet Alcedinidae purples are less saturated and have lower UV content than those in Sturnidae (Fig. S2b,c). Icteridae shows innovation in the red parts of color space (Fig. 2d). Furnariida shows little color novelty. This clade uses almost exclusively melanin pigments as color-producing mechanisms, which is also commonly seen in the other clades (Fig. 2e; Table 1).

**Table 1.** Suite of color-producing mechanisms varies by avian clade. Plumage color-producing mechanism data obtained from both general^13–16, 36, 78^ and clade-specific sources^19, 20, 28, 43, 79^. Dashes indicate a mechanism is absent in that clade. Note: number of species refers to those in our data set, not total species richness for a clade.

| Mechanism | Clade / no. species |  |  |  |  |
| --- | --- | --- | --- | --- | --- |
|  | Thraupidae<br>n=327 | Sturnidae<br>n=45 | Alcedinidae<br>n=72 | Icteridae<br>n=87 | Furnariida<br>n=604 |
| Carotenoids |  |  |  |  |  |
| Red carotenoids | common | – | – | common | – |
| Orange carotenoids | (rare) | – | (rare) | common | – |
| Yellow carotenoids | common | (rare) | (rare) | common | (rare) |
| Melanin |  |  |  |  |  |
| Eumelanin | common | common | common | common | common |
| Phaeomelanin | common | common | common | common | common |
| Structural coloration |  |  |  |  |  |
| Structural barb rami | common | – | common | (rare) | – |
| Structural barbule | (rare) | common | (rare) | common | – |
| White | common | common | common | common | common |
| Total mechanisms: | 8 | 5 | 7 | 8 | 4 |
| Mechanism score:<br>(absent = 0, rare = 1,<br>common = 2) | 14 | 9 | 11 | 15 | 7 |

### Evolutionary rates differ among clades and among patches

To understand whether clades and feather patches are evolving at different rates, we compared rates for each patch separately for males and females using a multivariate phylogenetic comparative approach^38^. We found significant rate variation among both clades and patches (both p < 0.01; Fig. 3b,c). Evolutionary rates were significantly elevated in male Thraupidae, Icteridae and Sturnidae, but in Furnariida females had significantly faster rates and in Alcedinidae rates were similar between males and females (Fig. 3b, Table S3). Furnariida showed the slowest evolutionary rates for both males and females (Fig. 3b). Comparing across clades, evolutionary rates of female coloration were significantly elevated in Alcedinidae compared to Furnariida, Thraupidae, and Sturnidae, whereas male rates were the same rate across all clades but Furnariida (Fig. 3b). Across all clades, the most rapid color evolution was seen in the belly and breast (Fig. 3c), a pattern driven primarily by Icteridae and Alcedinidae (Fig. S4). However, this pattern was not driven only by bright, colorful plumage patterns when considering the overall drabber colors of Icteridae relative to Alcedinidae (Fig. 2). Across all clades, the slowest pace of color evolution was found in the tail (Fig. 3c). An exception to this pattern was Sturnidae, with tail color rates 3.1 times faster than other clades (Fig. S4). Evolutionary rates of belly coloration in Sturnidae were significantly faster than other groups (Fig. S4). Alcedinidae was an outlier in terms of rates of dorsal (crown, rump and back) coloration, for both males (Fig. S4) and females (Fig. S5).

**Fig. 3.**
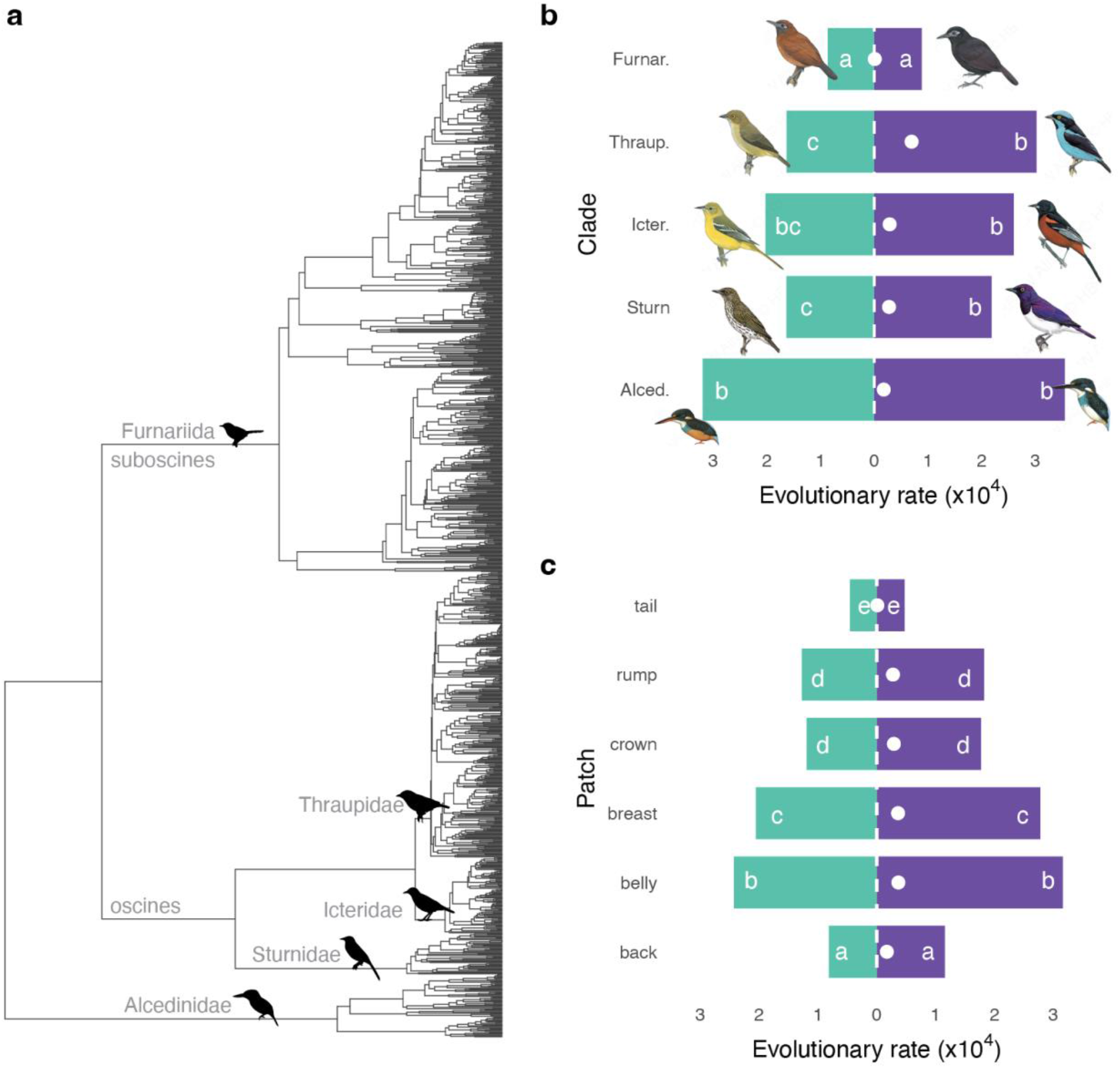
Evolutionary rate variation among clades and plumage patches. (a) Phylogenetic tree for five focal clades. Bar plots show multivariate evolutionary rates of coloration (UV XYZ coordinates and luminance) (b) among clades and (c) among plumage patches for males (purple) and females (green). Similarly colored bars sharing similar letters are not significantly different (p > 0.05). White circles indicate the relative rate difference between males and females. See Figs. S4, S5 for patch-wise rate analyses for each clade and sex.

### Clades with more color-producing mechanisms evolve color faster

To test whether the number of color-producing mechanisms available to a clade explains differences in the observed evolutionary rate variation (Fig. 3b), we used a phylomorphospace approach^39, 40^. Briefly, we determined the number of color mechanisms for each clade (Table 1), following an existing scoring terminology^16^, and then calculated the sum of color branch lengths for each clade (see Methods for details), a measure of disparity, and also the volume of an ellipsoid encompassing the points in 3-D color space^40^. We then calculated evolutionary rates as the sum of color branch lengths divided by the sum of branch lengths (in My) and lineage density as the sum of color branch lengths divided by the volume in color space. We compared these two metrics to the number of mechanisms, and mechanism scores, using standard regressions. This analysis revealed a positive relationship between the number of color-producing mechanisms and the evolutionary rate of a clade (Fig. S8c). Lineage density (Fig. S8d) and color space volumes (Fig. S8b) were not strongly associated with the number of color-producing mechanisms.

### Males and females have similar levels of plumage integration

To test the prediction that clades vary in the strength of covariation between color of different patches (i.e., plumage integration), we first estimated multivariate correlations (using paleomorph^41^) between each of the six focal patches in each clade. We did this for both the males and females data sets separately, using UV XYZ and luminance color variables. These comparative analyses revealed significant differences among clades in their plumage integration levels. For males, Furnariida showed significantly stronger integration than all clades except Icteridae (Fig. 4c). The three clades with diverse structural color mechanisms (Sturnidae, Thraupidae, Alcedinidae) showed significantly lower levels of integration than the primarily pigment-based clades Icteridae and Furnariida (Fig. 4c, Table 1). Similarly, in females, plumage integration was significantly elevated in Icteridae and Furnariidae relative to the other three clades (Fig. 4c). These cladewise differences in integration were related to evolutionary rate differences, with the fastest rates seen in clades with lower plumage integration (i.e., more modular plumages), but only in males (Fig. 4a).

**Fig. 4.**
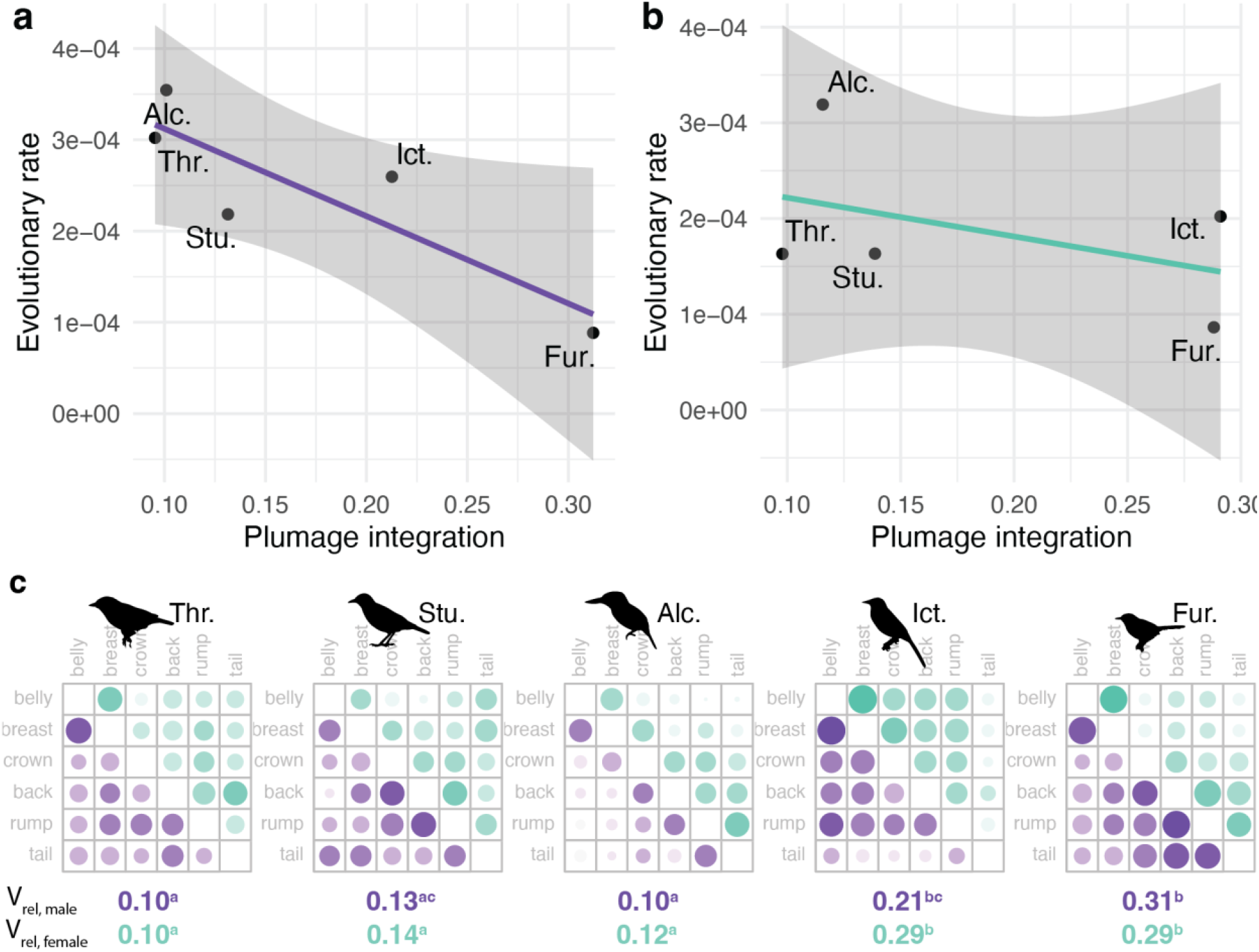
Relationship between plumage integration and evolutionary rates. Upper panels show relationship between plumage integration (V_rel_) and evolutionary rates for (a) males and (b) females. c, pairwise correlation plots for plumage patches (rows, columns) in each clade. Size of circles indicates strength of correlation between a given pair of patches. Color indicates males (purple) and females (green). Values below plots give the level of plumage integration (V_rel_) along with significance levels (numbers within a row sharing the same letter are not significantly different, FDR corrected P values). Differences between male and female plumage integration were not significant (V_rel_ effect size comparison, P > 0.05, Table S2).

### Interpatch color variation aligns with the direction of color evolution in females

To further test our prediction that variation among patches at the plumage level would explain evolutionary trends in color, we compared the major axes of phenotypic variation in color (P_max_) at both the plumage and clade levels. We used a Baysian phylogenetic mixed modeling (BPMM) approach^42^ to fit a multivariate response model (X, Y, Z, luminance variables) with phylogenetic covariance and patches as random effects. Using these flexible models, we were able to compare divergence in the major axes of interpatch and interspecific color variation. This angle was generally significantly different from zero in males, with the exception of Sturnidae (Table 2). By contrast, in females, the divergence angles were not significantly different except for Alcedinidae (Table 2, Fig. S9). Looking at these clades in color space, the major axis of male color divergence in Alcedinidae occurs in the XY plane (i.e., chromatic variation), with patches evolving along dark blue-light yellow colors and species diverging along a turquoise-darker phaeomelanin color axis (Fig. 5). In most clades showing a significant divergence between the major axes of interpatch and interspecific color variation, interpatch variation varies along a lightness axis while interspecific variation evolves along primarily chromatic axes (Fig. 5).

**Fig. 5.**
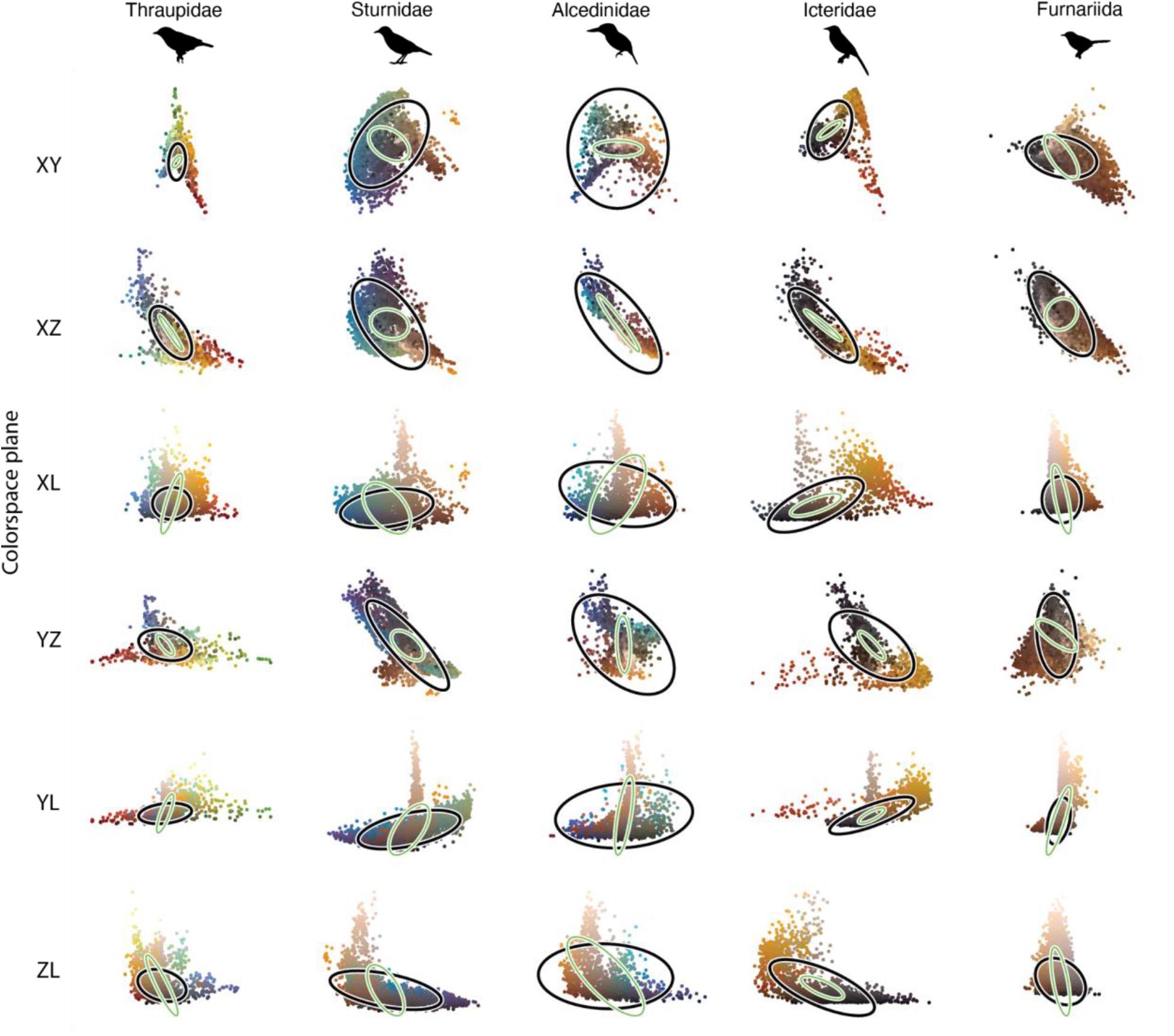
Directions of interpatch and interspecific color variation differ in males. Plots of 4-D tetrahedral color space for different planes (rows) and avian clades (columns). Ellipses show the major axes of color variation among species (black) and among patches on a bird’s body (green lines). Tetrahedral color space (TCS) coordinates were calculated using an ultraviolet-sensitive (UVS) visual system. Statistical analyses were also done using a violet-sensitive visual system (see Table S4). Note: axes were scaled by clade to aid in comparison of trends along a column. See Fig. S9 for results for female coloration.

**Table 2.** Divergence in major axes of color variation among patches and species. Significant results indicated in bold. Visual model assumes a UV-sensitive visual system with luminance calculated as the quantum catch of a blue tit visual system double cone^80^. 𝜃: angle between the major axes of interpatch and interspecies color variation. See Fig. 5, ref. ^63^ for further methodological details.

| Clade | Sex | Mean $\theta$ (°) | 95% credible interval | P-value |
| --- | --- | --- | --- | --- |
| **Thraupidae** | **Male** | **61.6** | **[40.1, 78.5]** | **0.02** |
| Thraupidae | Female | 68.0 | [40.9, 120.0] | 0.10 |
| Sturnidae | Male | 95.6 | [26.2, 140.3] | 0.08 |
| Sturnidae | Female | 113.1 | [29.8, 152.2] | 0.10 |
| **Alcedinidae** | **Male** | **49.1** | **[32.0, 64.7]** | **<0.01** |
| **Alcedinidae** | **Female** | **42.8** | **[29.7, 54.4]** | **<0.01** |
| **Icteridae** | **Male** | **16.7** | **[5.5, 28.4]** | **0.02** |
| Icteridae | Female | 11.8 | [4.5, 19.9] | 0.09 |
| **Furnariida** | **Male** | **109.5** | **[79.1, 135.4]** | **0.03** |
| Furnariida | Female | 95.4 | [50.8, 126.3] | 0.17 |

## Discussion

We tested the hypothesis that the major axes of interpatch and interspecies color variation are aligned within each of five phylogenetically diverse clades of birds. We found strong support for the prediction that plumage integration and rates of color evolution vary among clades with different color-producing mechanisms, yet these differences were dependent on whether we analyzed males or females. Color evolution occurred perpendicular to the axis of interpatch variation in males, but not in females. Stronger levels of plumage integration were further associated with slower rates of color evolution. This suggests that evolving new ways of producing color and flexibility at the developmental level (i.e. the ability to produce different colors across the body) are key factors in broad phylogenetic trends of coloration.

Novel plumage colors can be explained either by the evolution of new color-producing mechanisms, or by species evolving new colors with existing mechanisms. A comparison of color space volumes among groups of birds with different ways of producing colors led to the hypothesis that structural colors have expanded the avian color space^16^. Consistent with this idea, the largest color space volumes are seen in clades that commonly deploy structural coloration in their plumages (Fig. 2a-c). By contrast, the only clade without known structural color (Furnariida) occupied the lowest amount of color space (Fig. 2e). Convergence in color space between Alcedinidae and Sturnidae (Fig. 2b,c) is likely due to these clades having the greatest similarity in color-producing mechanisms (primarily structural colors; see Table 1). However, this pattern is interesting because each clade produces different forms of structural coloration in different parts of the feather. Whereas African starlings display a rainbow of iridescent colors emanating from melanin structures in feather barbules^37^, kingfishers produce non-iridescent (i.e., angle-independent) turquoise and blue structural colors through organized keratin structures in feather barb rami^43^. This result provides an example of how convergent phenotypes can result from divergent physical (or genetic) mechanisms, emphasizing the importance of identifying the specific mechanisms of color production^44^. Another way that birds can produce more colors from the same set of pigments is by using modified feather structures in combination with those pigments. For example, the carotenoid red colors of Thraupidae occur in a distinct part of color space compared to likely carotenoid red colors in Icteridae^45^ (Fig. 1a,d). This is possible either through a divergence in carotenoid types between these clades, or due to the interaction between feather microstructure and pigments in tanagers^46^. Finescale morphological diversity (e.g., dimensions of melanin structures in barbules) or interactions between pigments and feather structures could potentially explain the lack of an association between the number of color-producing mechanisms and color space volume (Fig. S8b). However, another possibility is that expansion in color space is driven by evolutionary shifts in rates of color evolution for different color-0producing mechanisms.

Clade comparisons are a powerful tool for understanding why some groups evolve more rapidly than others. The three clades with the greatest number of color mechanisms–Thraupidae, Icteridae, and Alcedinidae–shared similarly rapid rates of color evolution (Fig. 3b). The Sturnidae clade was also indistinguishable from these clades in terms of evolutionary rate of male coloration (Fig. 3b), despite a limited suite of color-producing mechanisms (Table 1). However, Sturnidae have iridescent structural coloration^28, 37^ for which a broader range of hues are possible whereas Alcedinidae and Thraupidae primarily utilize non-iridescent structural coloration^13, 36, 43^. Thus, it could be that iridescent colors evolve faster than non-iridescent structural colors. Another possible explanation is cryptic morphological diversity in feather nanostructures that is not captured in broad classes of color-producing mechanisms (Table 1). African starlings are known to produce iridescent color using a diverse set of melanin nanostructures found in feather barbules^28^, whereas non-iridescent structural color in Alcedinidae and Thraupidae may stem from more highly conserved morphotypes in the barb ramus^13, 47^. Testing this hypothesis will require more detailed microscopic information from across these non-iridescent radiations. Given the previously suggested hypothesis that clades with structural coloration expand their color space relative to clades with pigment-based coloration^16^, the rapid rates of color evolution in Icteridae (a clade known to display vivid carotenoid-based colors) are intriguing (Fig. 3b). Per-wavelength rate contours revealed that Icteridae males and females evolve rapidly in the yellow-red part of the spectrum (Figs. S6d, S7d), and the shapes of these rate contours are remarkably similar to those of carotenoid-based reflectance spectra^48^. This suggests that the mechanism for the high rates in Icteridae is rapid switching between melanin- and carotenoid-based coloration within patches. By contrast, Alcedinidae, Thraupidae, and Sturnidae show more even rate contours (Fig. S6a-c), consistent with the idea that color evolution proceeds primarily through variation in color produced by a single mechanism (e.g., iridescent structural color) or, alternatively, many switches among different mechanisms flattening out the rate contour. Fewer color-producing mechanisms in Furnariida (Table 1), along with higher plumage integration (Fig. 4c), likely both contribute to the slow evolutionary rates of color in this clade (Fig. 3b). Interestingly, color disparity was highest in female Furnariida (Fig. S8a), yet high lineage densities in this clade (Fig. S8c) are indicative of tight clustering in color space (Fig. 2e) and slow rates of evolution to other parts of color space (Fig. 3b). Across all clades, the lack of a trend between lineage density and number of color-producing mechanisms (Fig. S8d) suggests that rather than species jumping to new parts of color space by evolving novel color-producing mechanisms^16^, species are evolving faster for a given set of mechanisms, in line with recent work on the evolvability of iridescent feather nanostructures^29, 30^.

Sexually selected traits evolve more rapidly than naturally selected traits^49^, therefore comparing evolutionary dynamics of coloration between sexes and among plumage patches can inform us about relative roles of sexual and natural selection in diversification^50–53^. A recent study in the avian Tyrannida clade found that rates of color evolution were faster in more sexually dichromatic species^54^. This study used lineage-specific evolutionary rates, whereas we use clade-based analyses here, but lineage-specific rates found significant correlations between diversification rates and rates of color evolution in both male and female tanagers^55^. At the clade level, we find similar and rapid evolutionary rates of male coloration in both a dichromatic clade^52^ (Thraupidae), in a monochromatic clade^51^ (Alcedinidae; Fig. 3b), and in a clade that includes both di- and monochromatic linages (Furnariida). One possible explanation for this difference is that males and females have more similar plumage integration levels in these clades, whereas in other dichromatic clades male plumage patches are more decoupled than in female plumage patches. However, levels of plumage integration did not differ significantly between males and females (Fig. 4c). Consistent with previous work^54^, we also found that evolutionary rates of coloration were generally fastest in crown and breast patches and slowest in the tail (Fig. 3b), but some dorsal regions also showed elevated rates, specifically the rump patch (Fig. 3c). This pattern was driven primarily by the Alcedinidae clade (Figs. S4, S5) in the UV and blue parts of the spectrum (Fig. S7c). Rump coloration would be visible to conspecifics during spread-wing courtship displays characteristic for the group^56^. If elevated rates in dorsal patches in Alcedinidae imply differences in selective regimes between dorsal and ventral parts of the body^57, 58^ (e.g., sexual selection for dorsal patches versus natural selection on ventral patches), this could weaken developmental links between body regions^59^, thereby reducing evolutionary covariation among color patches. Yet, we did not find support for this idea, as plumage integration levels were similarly low in Sturnidae, and Thraupidae compared to Alcedinidae (Fig. 4c). Selection on alternative (i.e., non-signaling) functions of plumages might also be driving some of the observed clade-specific patterns (Fig. 3). For example, rapid rates of tail color evolution in Alcedinidae and Sturnidae–two clades known to produce structural coloration with melanin pigments–could allow these clades to elaborate their plumage colors while at the same time maintain a wear-resistant or mechanical function in stabilization during flight or climbing^44, 60^. In other clades with primarily pigment-based mechanisms (e.g., Icteridae and Furnariida), switches to carotenoid coloration in the tail could lead to increased wear of feathers due to abrasion^61, 62^.

Strong covariation among plumage patches can limit or enhance evolutionary diversification of color, depending on the alignment of the axis of variation relative to that of selection^1^. At the overall plumage level, faster rates of male color evolution were associated with low plumage integration levels (Fig. 4a). This suggests that more modular plumages promote color diversity in males, echoing results at the single patch level showing that low levels of integration between nanostructural subtraits promotes rapid rates of color evolution^29^. A recent study found that species with more distinct plumage patches evolve overall plumage coloration at a faster rate than species with fewer patches^32^, but this study did not consider plumage integration between patches as we do here. Compared to the low levels of plumage integration in Alcedinidae, Thraupidae, and Sturnidae, high levels of plumage integration in Icteridae and Furnariida females (Fig. 4c) are consistent with stronger developmental constraints in these clades. Yet, strong covariation among color patches at the plumage level (i.e., a narrow ellipse in color space) does not necessarily imply a similar pattern at evolutionary scales, as we found significant support for divergence in the directions of interpatch and interspecific color variation in most clades (Fig. 5). The direction of interpatch divergence was generally along the luminance axis (i.e., plumage lightness), whereas evolutionary change proceeded more along chromatic axes (Fig. 5). This makes sense if color mechanisms are more easily lost among patches (e.g., structural blue to white, or carotenoid orange to white transitions) than among species, possibly owing to a shared developmental toolkit influencing where on the body colors can be turned “on” or “off”^22^. One exception to interpatch variation along the luminance axis was Icteridae, as the interpatch axis was more closely aligned to the interspecies black-red/yellow color axis (Fig. 5), suggestive of flexible switching between carotenoid and non-carotenoid coloration at both plumage and evolutionary scales. Previous studies have compared the alignment of the major axis of phenotypic variation (P_max_) at intraspecific^63^ and interspecific levels^64^, but to our knowledge none have compared P_max_ across scales. Taken together, divergence between interpatch and interspecies P_max_ vectors suggests that plumage patterns constrain evolution of color in these clades, whereas, in clades with similarly aligned major color axes, there is less constraint on the direction of color evolution, and color is free to evolve along “lines of least resistance”^1^.

## Conclusions

Our spectral data set covers nearly all pigment types^14, 15^ and color-producing feather nanostructures^13, 65^ known in birds (Table 1), with the exception of some more uncommon pigments (psittacofulvin, turacoverdin). Compared to recent work on color evolution using alternative data sources (e.g., color plates^50^ and digital photography^54, 66^), reflectance spectra have enabled us to look at per-wavelength evolutionary rates (Figs. S6, S7) and make richer inferences about color mechanisms and evolution compared to using color space coordinates alone. We hope that future work will continue to assess how different ways of quantifying plumage coloration might lead to similar (or different) conclusions^67^. Our finding that color evolution across species has proceeded in a different direction than among color patches at the individual species’ plumage level (Fig. 5) shows how plumage color patterns can be an ideal system for exploring the interplay between innovation and constraint in driving phenotypic evolutionary trends.

## Methods

### Phylogenies

We obtained published time-calibrated phylogenies for Icteridae^68^ and Thraupidae^69^, Alcedinidae^70^, Sturnidae^28^, and Furnariida^71^. We then combined these subtrees into a larger supertree using the bind.tree function in ape^72^ based on published divergence times between the clades (http://www.timetree.org).

### Measuring feather reflectance

We used an Ocean Optics USB2000 spectrophotometer (Dunedin, FL) with a PX-2 pulsed xenon light source to record reflectance across the avian visual spectrum, ranging from 300 to 700 nm. We used a R200-7-UV/VIS reflection probe fitted with a modified rubber stopper to restrict incident light and to control the distance between the probe tip and feather surface (∼1 cm). All measurements were taken with the light and probe perpendicular to the feather surface (i.e. normal incidence). The number of patches measured varied for each clade (Thraupidae n = 9, Sturnidae n = 13, Alcedinidae n = 22, Icteridae n = 20, Furnariida N = 8), with six patches in common to all data sets (Fig. S1). We took three replicate measurements of each region for each individual and up to 11 individuals per sex per species and averaged them for subsequent analyses. After removing data for species not in the phylogeny, our final spectral data set contained 45791 mean reflectance spectra (n = 5886 Thraupidae, n = 7310 Sturnidae, n = 3079 Alcedinidae, n = 5362 Icteridae, n = 24154 Furnariida) across 1135 species (n = 327 Thraupidae, n = 45 Sturnidae, n = 72 Alcedinidae, n = 87 Icteridae, n = 604 Furnariida).

### Avian visual models

We performed all visual model analyses in the R package pavo^35^. Briefly, we first ran avian visual models considering both a UV-sensitive (UVS) and violet-sensitive (VS) visual system using the vismodel function. We next calculated tetrahedral color space (TCS) coordinates for each spectrum using colspace, resulting in two data sets (UVS and VS) of XYZ coordinates. To assess how robust our analyses are to assumptions about the visual capabilities of our focal clades, we performed comparative analyses using both UVS and VS visual systems.

### Estimating evolutionary rates

Using color space coordinates, we transformed data into a species by patch-coordinate matrix (e.g., *Ceyx margarethae* | wing-X wing-Y wing-Z…). We then estimated multivariate phylogenetic signal (Pagel’s 𝜆) using mvgls^73^ and transformed branch lengths according to the optimal 𝜆 value, for both males and females. Using geomorph^74^, we then compared evolutionary rates among clades with compare.evol.rates^38^ and among patches with compare.multi.evol.rates^75^. We estimated the significance of these relationships using a permutation approach (n = 999 simulations). Using these same methods, we also calculated rates using ln-transformed reflectance values in 20-nm bins.

### Comparing levels of plumage integration

To compare evolutionary covariation among patches (i.e., plumage integration), we calculated the V_rel_ metric in the geomorph function integration.Vrel for each clade and sex. We then compared V_rel_ values using the compare.ZVrel function^76^. We adjusted P-values for multiple comparisons with the false discovery rate (FDR) metric and calculated significance letters for each group using the multcompLetters function in the multcompView R package.

### Comparing major axes of color variation

To test our prediction that interpatch and interspecies color variation would differ in direction, we used Bayesian phylogenetic mixed models (BPMMs) implemented in the MCMCglmm R package^42^. We chose BPMMs over standard phylogenetic generalized least squares (PGLS) approaches because BPMMs are more flexible in that they allowed us to account for phylogenetic relationships and intraspecific measurements (e.g., spectral measurements on different patches of the same birds) in the same analysis. For the phylogeny, we used the merged supertree with untransformed branch lengths and estimated phylogenetic signal from the posterior variance-covariance estimates^77^. We used XYZ coordinates (plus luminance) as the multivariate response and patch and species as random effects. Using the posterior distribution of interpatch and interspecies covariance matrices, we then calculated the angle of divergence 𝜃 along major axes of color variation (P_max_) for patches and species in 3-D space (see Dryad for R code). To determine statistical significance of 𝜃, we followed a previous approach used to study genetic architecture in crickets^63^. Briefly, we subsampled 500 posterior MCMC samples, determined within and among group variation in 𝜃, and then calculated the test statistic 𝜙 as the total among-group 𝜃 variation minus the total within-group 𝜃 variation. We then calculated P values as one minus the proportion of 𝜙 values greater than zero.

