## Supporting Information for "Constraint and innovation in color evolution among species and among plumage patches in five avian radiations"

Author affiliations:
[1]Grainger Bioinformatics Center, Field Museum of Natural History, Chicago, IL 60605
[2]BioSciences Department, Rice University, Houston, TX 77005
[3]Museum of Natural Science and Department of Biological Sciences, Louisiana State University, Baton Rouge, Louisiana, 70803
[4]Biology Department, Drake University, Des Moines, IA 50311
[5]Data Scientist, Apple, Austin, TX
[6]Department of Biology, San Diego State University, San Diego, CA 92182
[7]Ornithology Department, Natural History Museum of Los Angeles, Los Angeles, CA 90007

Corresponding author: Chad M. Eliason


Keywords: evolutionary rates, spectrophotometry, iridescence, phenotypic evolution

*Supplementary Tables*

**Table S1. Color space volumes and overlap.** Color volumes on the diagonal (orange), raw overlaps are on the lower off-diagonals (blue), and proportional overlaps are in the upper off-diagonals (green). Results are shown for XYZ color space coordinates assuming a UV-sensitive visual system.

| | Clade | | | | |
|---|---|---|---|---|---|
| | Thraupidae | Sturnidae | Alcedinidae | Icteridae | Furnariida |
| Thraupidae | 0.066 | 0.313 | 0.214 | 0.227 | 0.134 |
| Sturnidae | 0.021 | 0.021 | 0.426 | 0.368 | 0.239 |
| Alcedinidae | 0.014 | 0.011 | 0.015 | 0.408 | 0.213 |
| Icteridae | 0.015 | 0.01 | 0.009 | 0.016 | 0.189 |
| Furnariida | 0.009 | 0.006 | 0.005 | 0.004 | 0.012 |

**Table S2. Plumage integration does not differ significantly between males and females.** Results of pairwise Z score comparison using compare.ZVrel in geomorph.

| **Clade** | **Vrel, male** | **Vrel, female** | **Pairwise Z** | **P** |
|---|---|---|---|---|
| Thraupidae | 0.095 | 0.098 | 0.18 | 0.85 |
| Sturnidae | 0.131 | 0.139 | 0.14 | 0.89 |
| Furnariida | 0.312 | 0.288 | 0.94 | 0.35 |
| Alcedinidae | 0.101 | 0.116 | 0.42 | 0.67 |
| Icteridae | 0.213 | 0.291 | 1.14 | 0.26 |

**Table S3. Color evolutionary rate difference between males and females.**

| Clade | Male rate | Female rate | Eff. size | P |
|---|---|---|---|---|
| Alcedinidae | 2.59E-04 | 2.87E-04 | 1.849 | 0.054 |
| **Furnariida** | **9.08E-05** | **8.51E-05** | **4.211** | **0.001** |
| **Icteridae** | **5.11E-04** | **6.41E-04** | **2.454** | **0.006** |
| **Thraupidae** | **3.73E-04** | **6.14E-04** | **17.220** | **0.001** |
| **Sturnidae** | **2.97E-04** | **3.62E-04** | **3.248** | **0.001** |

**Table S4. Divergence in major axes of color variation among patches and species.** Significant results indicated in bold. Visual model assumes a violet-sensitive visual system with luminance calculated as the quantum catch of a blue tit visual system double cone. $\theta$: angle between the major axes of interpatch and interspecies color variation.

| Clade | Sex | Mean $\theta$ (º) | 95% credible interval | P-value |
|---|---|---|---|---|
| Thraupidae | Male | 77.7 | [49.1, 125.1] | 0.27 |
| Thraupidae | Female | 93.6 | [47.3, 136.7] | 0.34 |
| **Sturnidae** | **Male** | **84.0** | **[26.0, 131.0]** | **0.03** |
| **Sturnidae** | **Female** | **98.7** | **[50.0, 143.7]** | **0.04** |
| **Alcedinidae** | **Male** | **53.7** | **[25.8, 92.6]** | **0.02** |
| **Alcedinidae** | **Female** | **46.7** | **[30.1, 62.7]** | **<0.01** |
| **Icteridae** | **Male** | **18.8** | **[4.2, 33.5]** | **0.02** |
| Icteridae | Female | 10.7 | [2.7, 19.6] | 0.16 |
| **Furnariida** | **Male** | **84.7** | **[52.7, 115.8]** | **0.02** |
| Furnariida | Female | 68.4 | [37.0, 104.9] | 0.07 |

*Supplementary Figures*

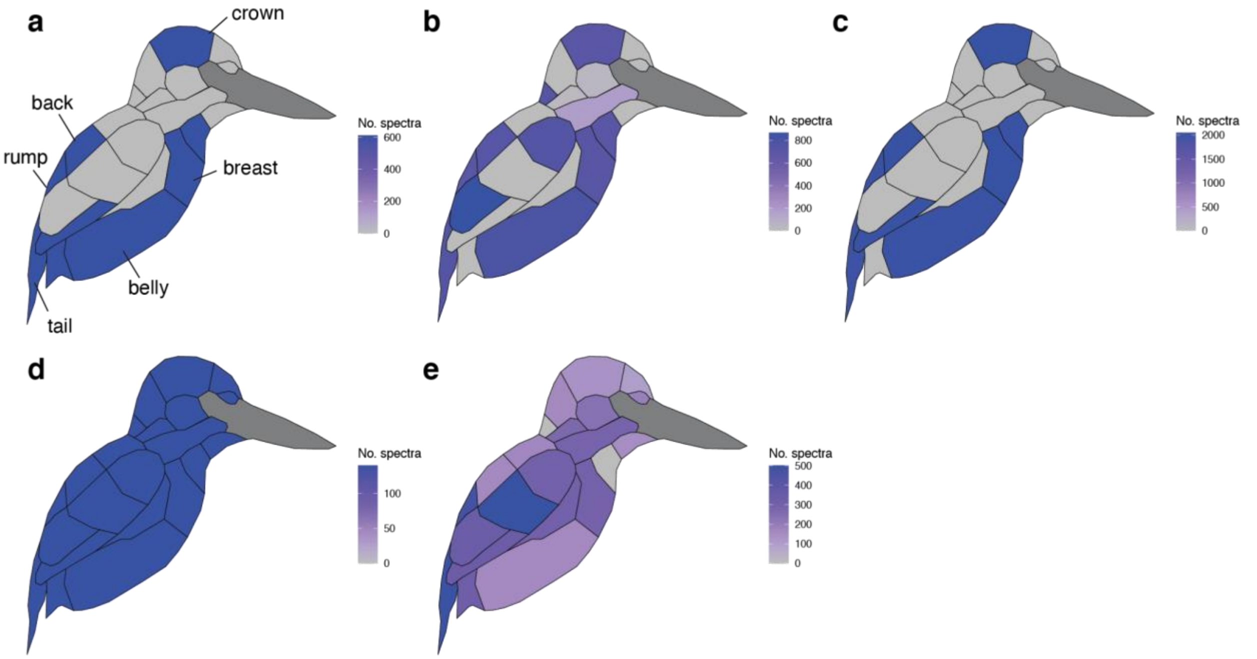


**Fig. S1. Number of spectra measured per body region.** Panels correspond to different clades: (**a**) Thraupidae, (**b**) Sturnidae, (**c**) Furnariida, (**d**) Alcedinidae, and (**e**) Icteridae. Six patches in common to all data sets are labeled in **a**.

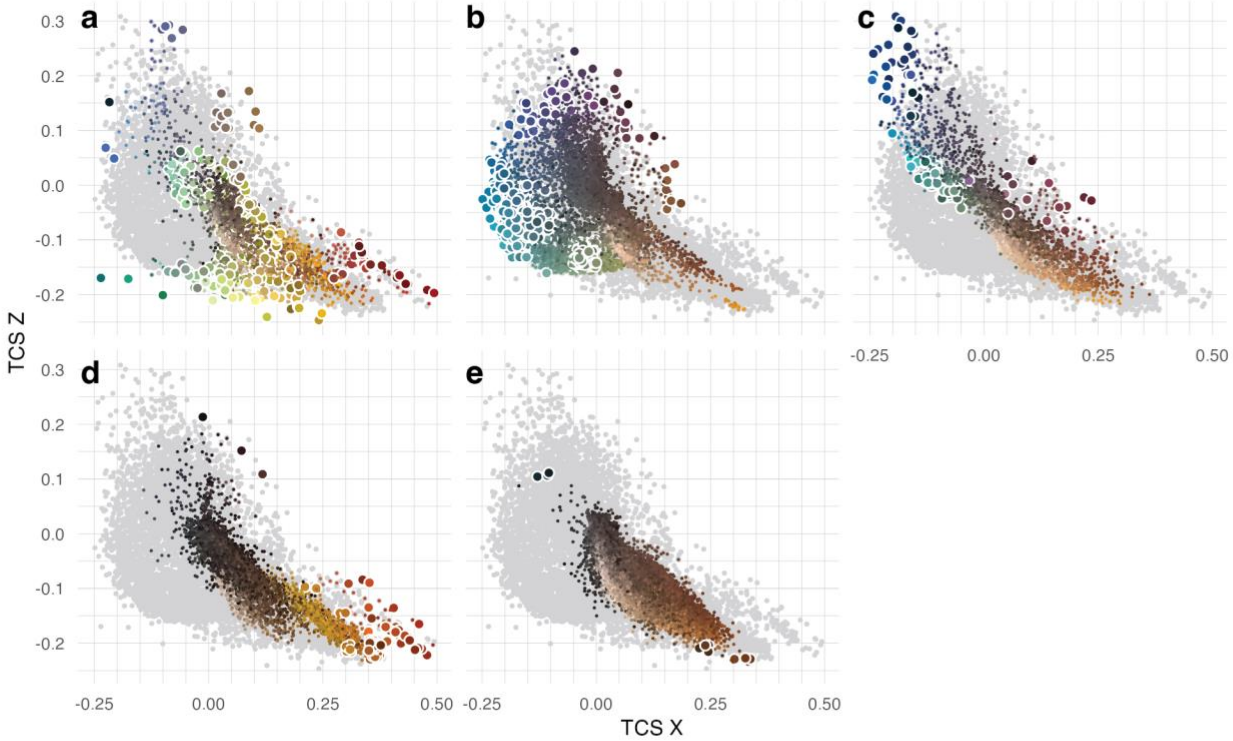


**Fig. S2.** Tetrahedral color space plots in the XZ plane for (**a**) Thraupidae, (**b**) Sturnidae, (**c**) Alcedinidae, (**d**) Icteridae, and (**e**) Furnariida. Gray points are all color space XY coordinates in our data set. Filled points are colors shared by two or more clades. Large points encircled in white depict novel colors in a clade. Lines delimit voxels dividing up color space.

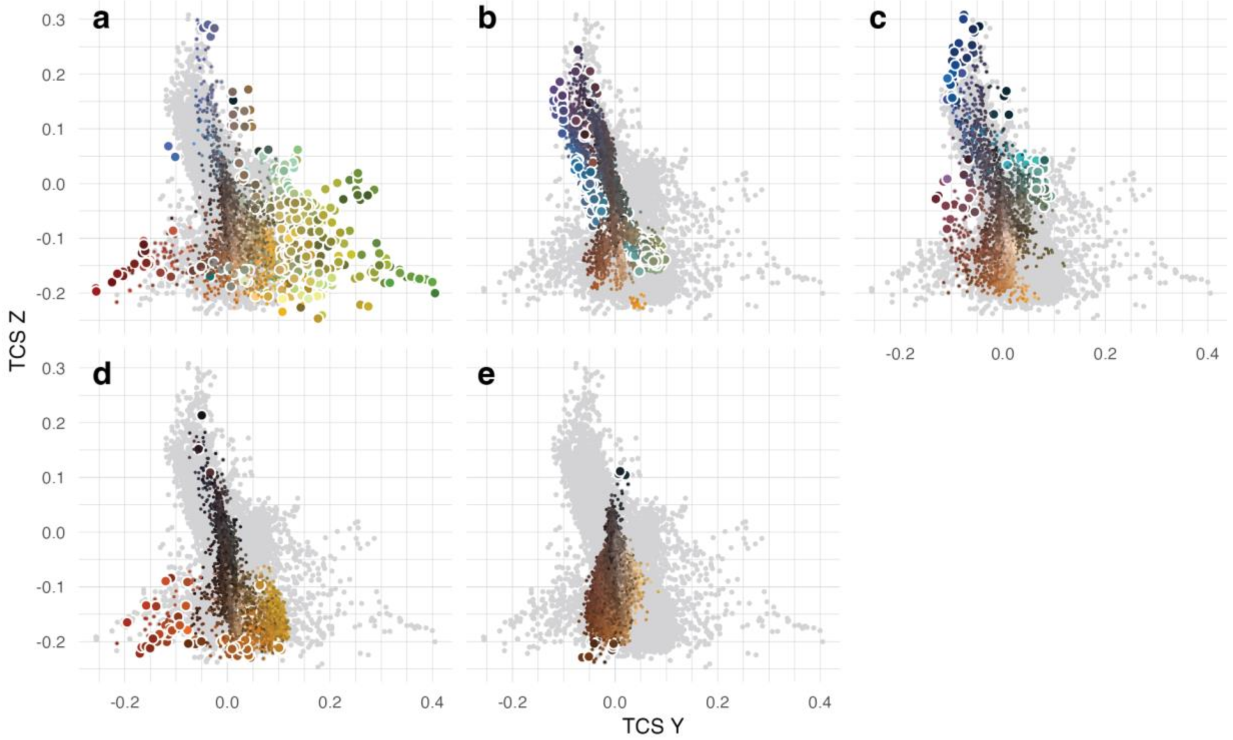


**Fig. S3.** Tetrahedral color space plots in the YZ plane for (**a**) Thraupidae, (**b**) Sturnidae, (**c**) Alcedinidae, (**d**) Icteridae, and (**e**) Furnariida. Gray points are all color space XY coordinates in our data set. Filled points are colors shared by two or more clades. Large points encircled in white depict novel colors in a clade. Lines delimit voxels dividing up color space.

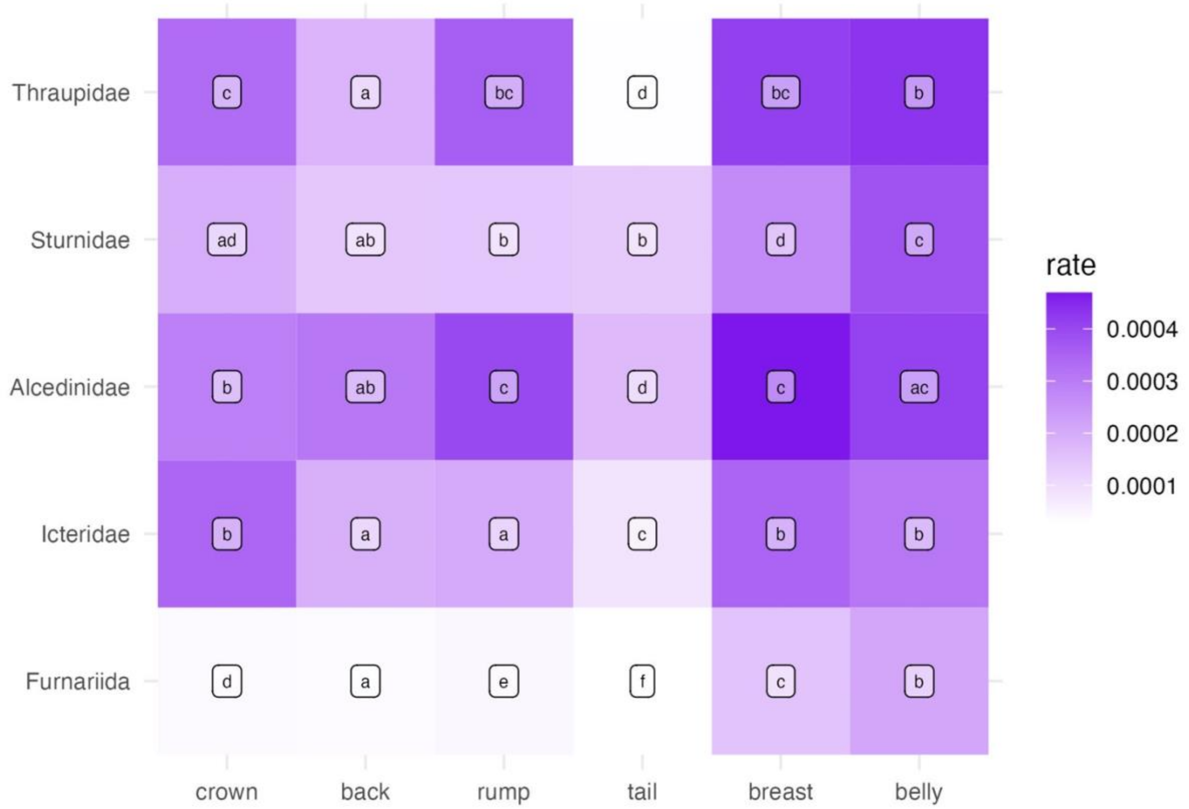


**Fig. S4. Rates of male color evolution by patch and clade.** Multivariate rates of evolution compared among patches (columns) and clades (rows) using compare.multi.evol.rates[75] in the geomorph R package[74]. Cell colors correspond to evolutionary rates (see legend). Cells sharing the same letter are not significantly different (P > 0.05). Note that statistical comparisons with letters are only valid within a row (i.e., clade). Rates shown for XYZ coordinates (plus luminance) derived from visual models assuming a UVS visual system.

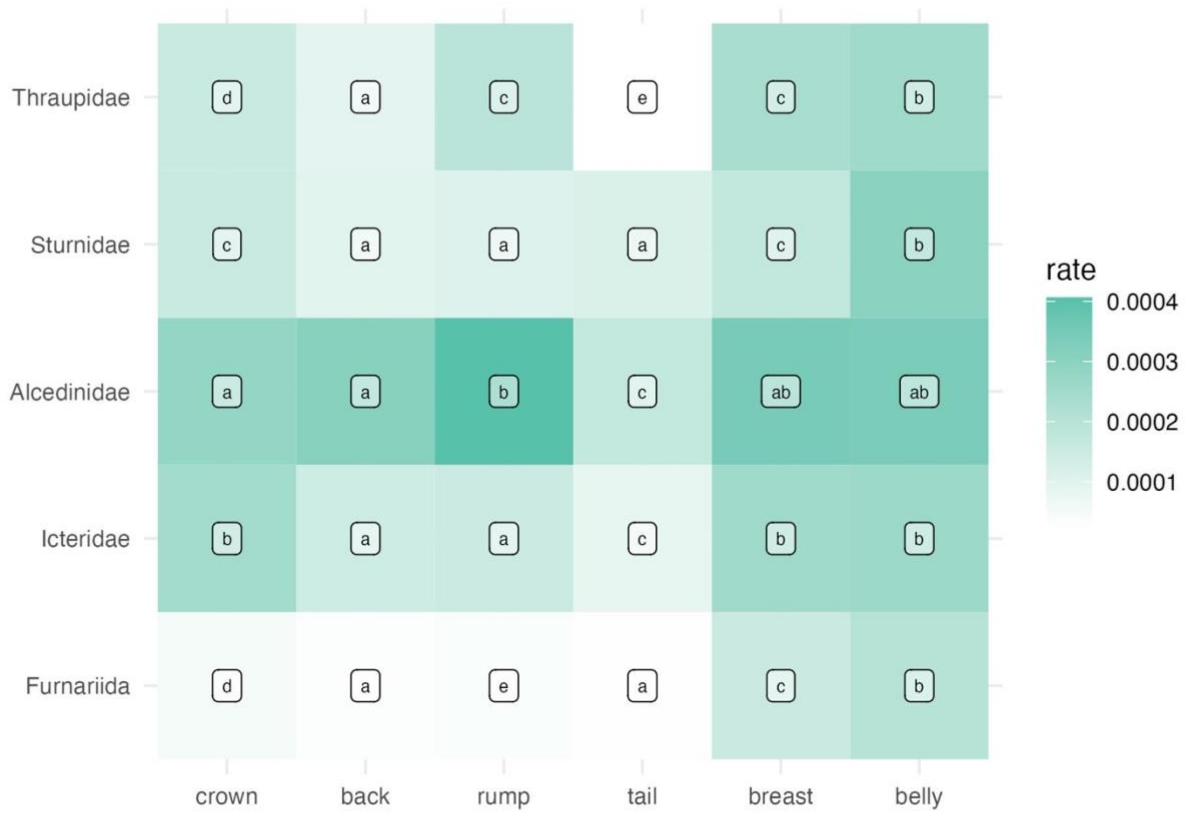


**Fig. S5. Rates of female color evolution by patch and clade.** Multivariate rates of evolution compared among patches (columns) and clades (rows) using compare.multi.evol.rates[75] in the geomorph R package[74]. Cell colors correspond to evolutionary rates (see legend). Cells sharing the same letter are not significantly different ($P > 0.05$). Note that statistical comparisons with letters are only valid within a row (i.e., clade). Rates shown for XYZ coordinates (plus luminance) derived from visual models assuming a UVS visual system.

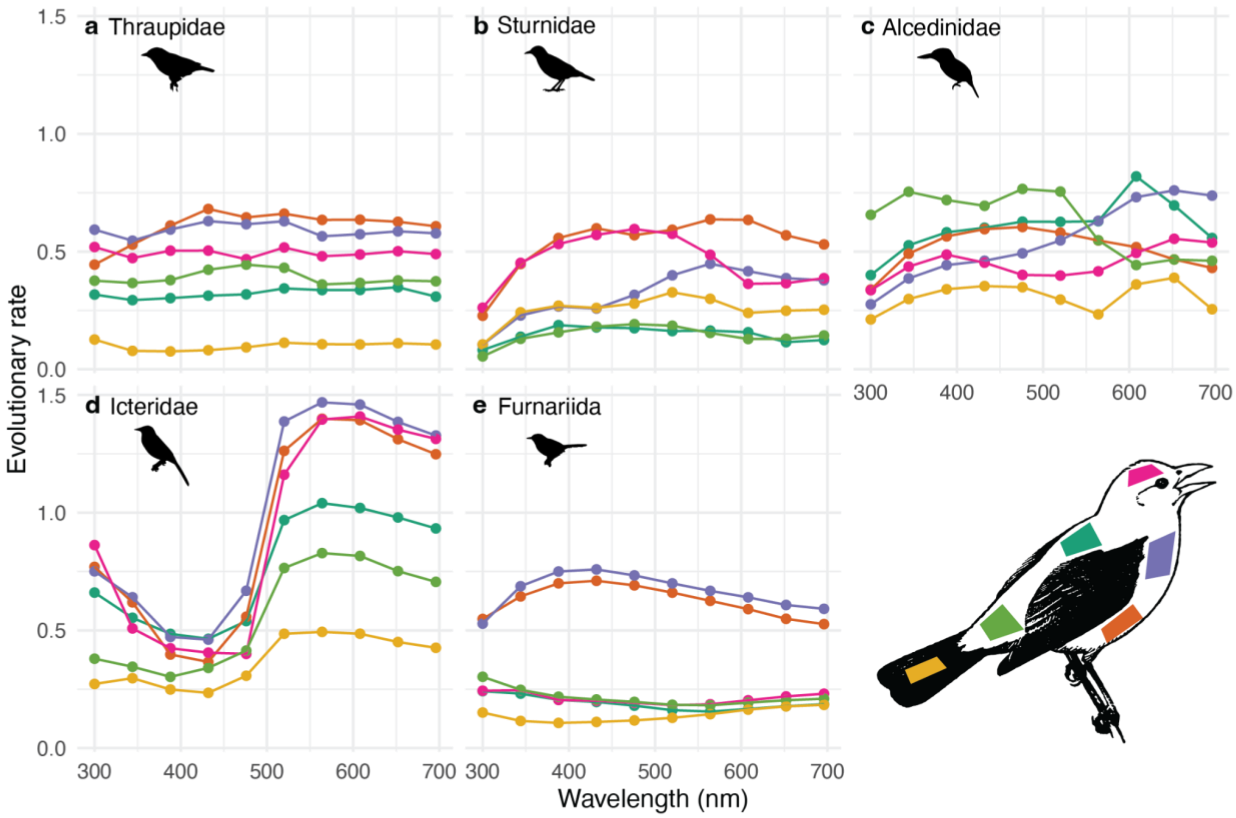


**Fig. S6. Per-wavelength evolutionary rates of male coloration for different clades and patches.** Reflectance values were ln-transformed prior to rate calculations with fit_t_pl in RPANDA. Plumage patches labeled in lower right. Image credit: scarlet tanager (*Piranga olivacea*) by Tirriko on pixabay (CC license).

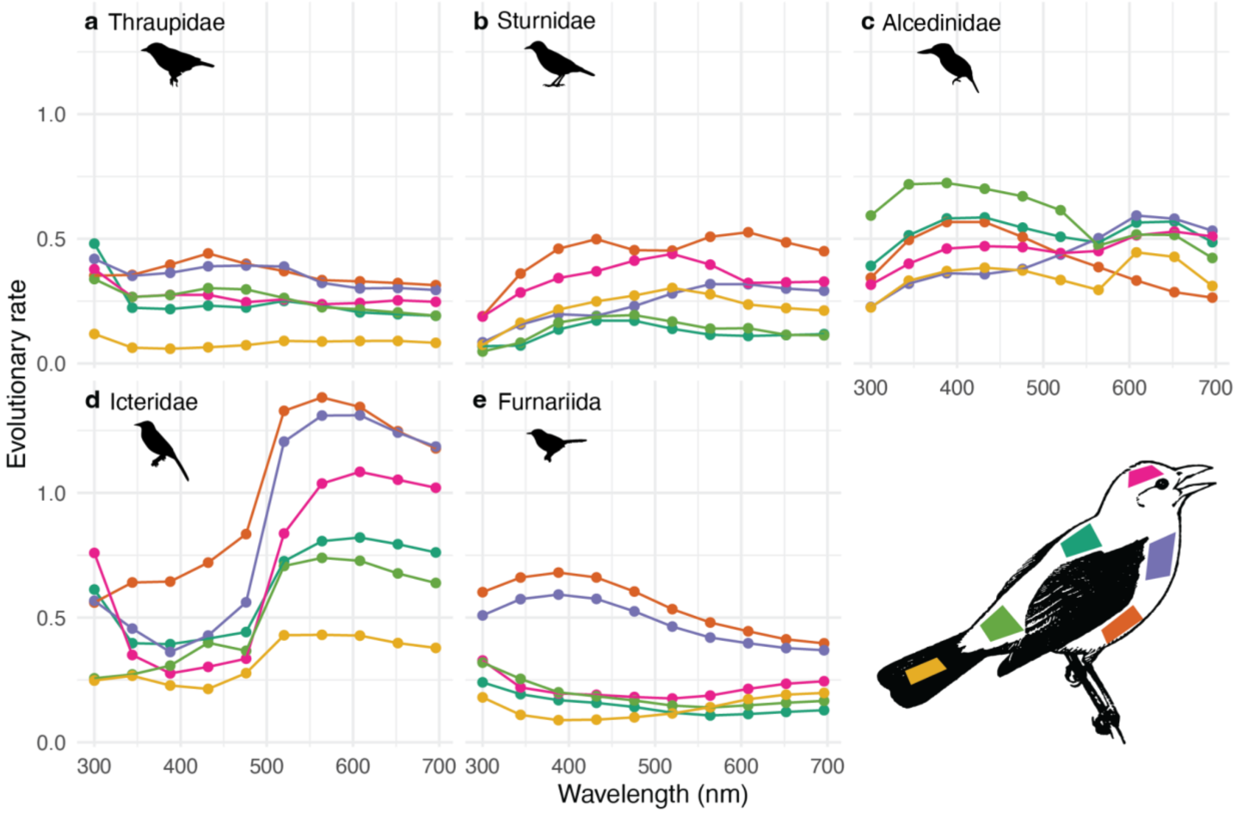


**Fig. S7. Per-wavelength evolutionary rates of female coloration for different clades and patches.** Reflectance values were ln-transformed prior to rate calculations with fit_t_pl in RPANDA. Plumage patches labeled in lower right. Image credit: scarlet tanager (*Piranga olivacea*) by Tirriko on pixabay (CC license).

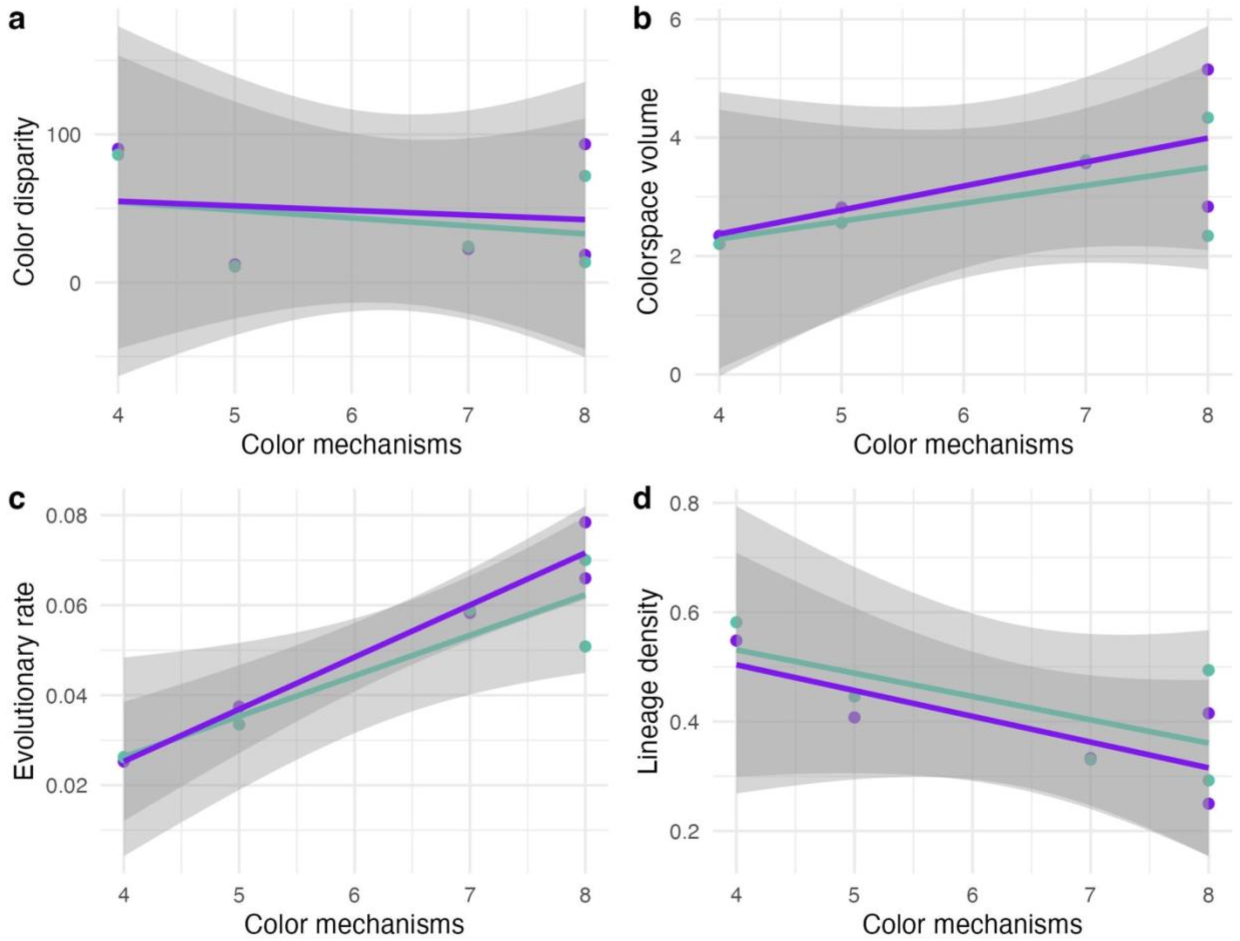


**Fig. S8. Color evolves faster in clades with more ways of producing feather coloration.** Phylomorphospace analysis showing color space disparity (**a**), color space volume (**b**), evolutionary rates (**c**) and lineage density (**d**) for males (purple) and females (turquoise). The positive association between the number of color-producing mechanisms and rate, but not lineage density, suggests that novel color-producing mechanisms enable clades to explore color space at faster rates.

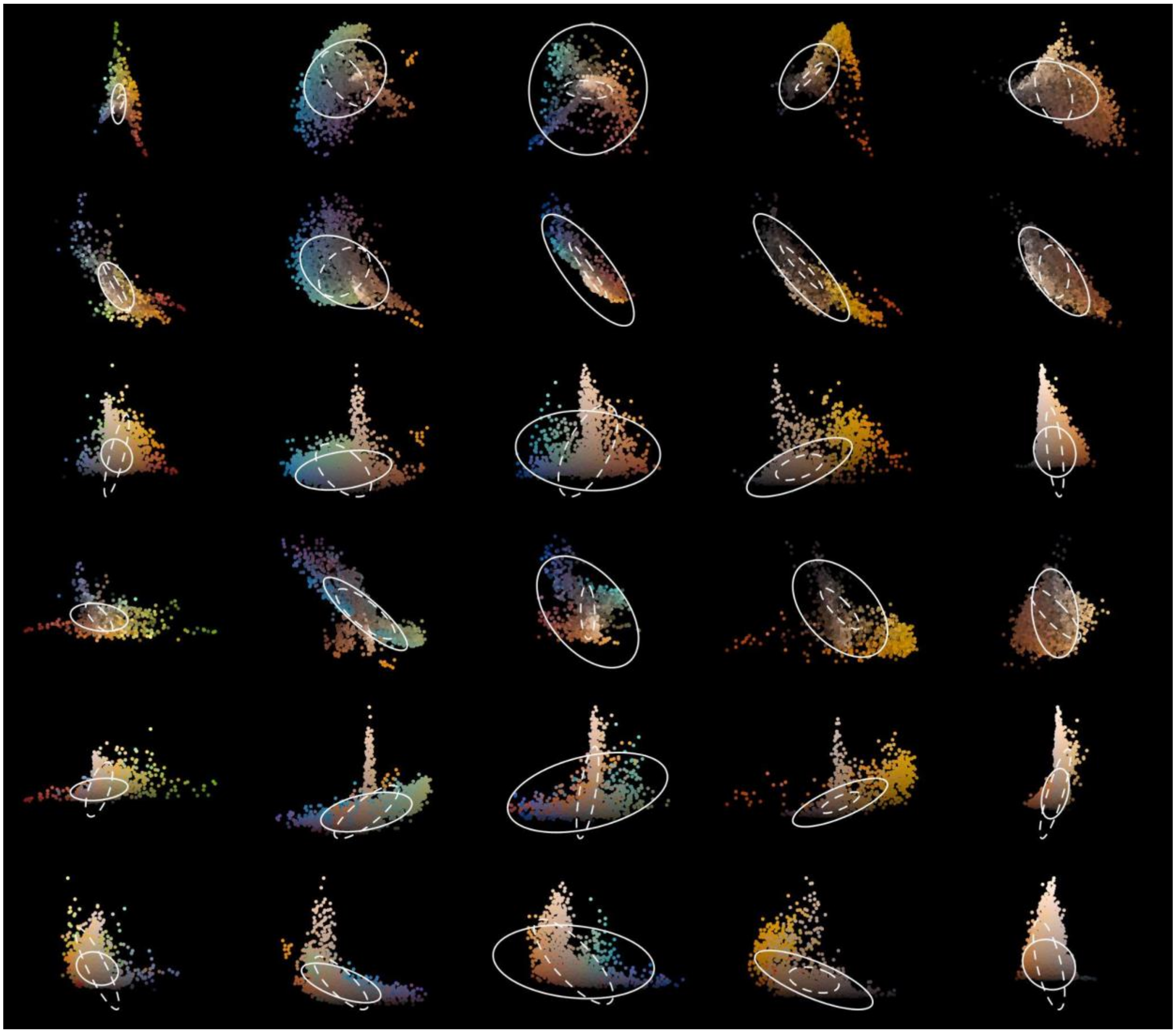

**Fig. S9. Color divergence among patches and species for females.** Panels show different planes of 3D color space (rows) for five focal clades (columns). Tetrahedral color space (TCS) coordinates calculated from visual models assuming a UV-sensitive visual system. Lines correspond to variation among patches (dashed) and among species (solid).

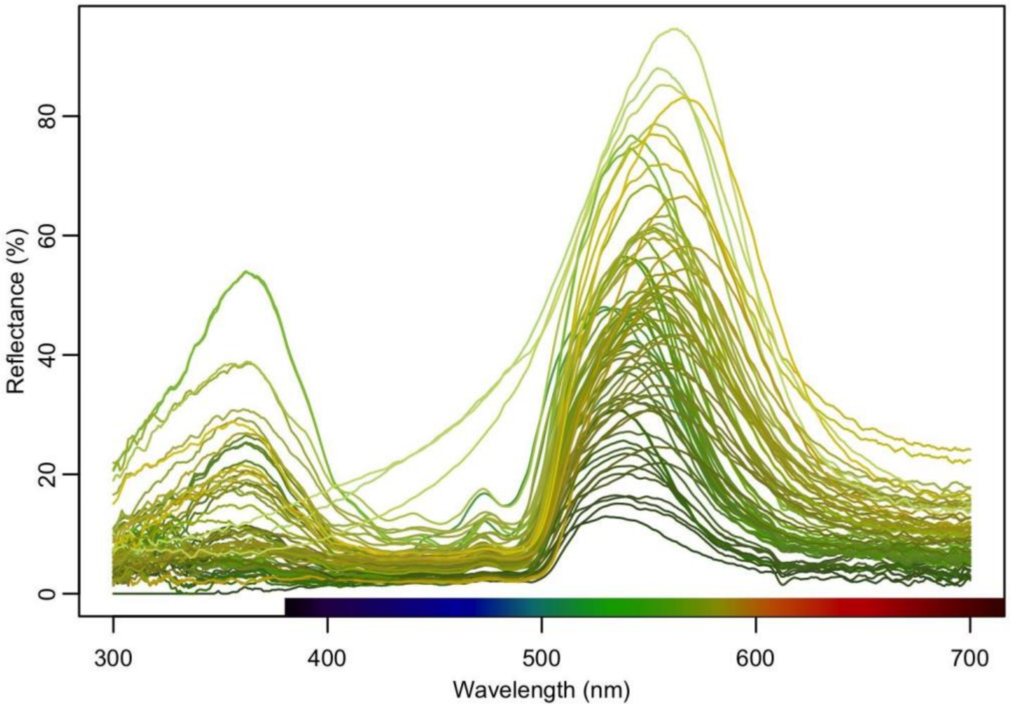


**Fig. S10. Reflectance spectra associated with novel yellow-green colors in Thraupidae.** See Fig. 1a for depiction of these colors in avian color space.
